# Sequence- and structure-selective mRNA m^5^C methylation by NSUN6 in animals

**DOI:** 10.1101/2020.10.03.324707

**Authors:** Jianheng Liu, Tao Huang, Yusen Zhang, Tianxuan Zhao, Xueni Zhao, Wanying Chen, Rui Zhang

**Author notes:** Correspondence (R.Z.), (T.H.). These authors contributed equally to this work (Jianheng Liu, Tao Huang, Yusen Zhang, Tianxuan Zhao).

## Abstract

mRNA m^5^C, which has recently been implicated in the regulation of mRNA mobility, metabolism, and translation, plays important regulatory roles in various biological events. Two types of m^5^C sites are found in mRNAs. Type I m^5^C sites, which contain a downstream G-rich triplet motif and are computationally predicted to locate in the 5’ end of putative hairpin structures, are methylated by NSUN2. Type II m^5^C sites contain a downstream UCCA motif and are computationally predicted to locate in the loops of putative hairpin structures. However, their biogenesis remains unknown. Here we identified NSUN6, a methyltransferase that is known to methylate C72 of tRNA^Thr^ and tRNA^Cys^, as an mRNA methyltransferase that targets Type II m^5^C sites. Combining the RNA secondary structure prediction, miCLIP, and results from a high-throughput mutagenesis analysis, we determined the RNA sequence and structural features governing the specificity of NSUN6-mediated mRNA methylation. Integrating these features into an NSUN6-RNA structural model, we identified an NSUN6 variant that largely loses tRNA methylation but retains mRNA methylation ability. Finally, we revealed a negative correlation between m^5^C methylation and translation efficiency. Our findings uncover that mRNA m^5^C is tightly controlled by an elaborate two-enzyme system, and the protein-RNA structure analysis strategy established may be applied to other RNA modification writers to distinguish the functions of different RNA substrates of a writer protein.

## Introduction

mRNAs contain numerous modified nucleotides, which have been suggested to play important roles in regulating mRNA fates. To date, how the writer proteins recognize specific sequence context for mRNA modification is largely unknown. Recently, it has been found that most writer proteins have multiple types of RNA substrates. For example, METTL16 is responsible for both snRNA and mRNA m^6^A modification [1], TRMT6/TRMT61A is responsible for both tRNA and mRNA m^1^A modification [2], and individual PUS proteins are responsible for tRNA and mRNA pseudouridylation [3]. Moreover, for all of these writer proteins, their mRNA substrates have similar features to tRNA or snRNA substrates [4]. These phenomena raise an intriguing hypothesis that, by mimicking sequence/structural features of noncoding RNAs, mRNAs co-opt those writer proteins to form modified nucleotides, although whether these modifications in mRNAs have become functionally adapted is still unknown.

RNA m^5^C is one of the longest-known RNA modifications. The presence and functions of m^5^C in noncoding RNAs, such as tRNA and rRNA, have been extensively investigated [5-11]. m^5^C has been recently found in mRNAs [12-14] and suggested to affect most posttranscriptional steps in gene expression [13, 15-17]. Functionally, mRNA m^5^C has been shown to impact various biological events [18-21], such as embryonic development, myelopoiesis, and cancer cell proliferation.

With the development of an experimental and computational framework to accurately identify mRNA m^5^C sites genome-wide, two types of mRNA m^5^C sites are found in animals [17]. Type I m^5^C sites are adjacent to a downstream G-rich triplet motif and predicted to locate in the 5’ end of stem-loop structures. Interestingly, NSUN2, a tRNA m^5^C methyltransferase, is responsible for Type I mRNA m^5^C methylation [12, 13], and the sequence and structural features of Type I mRNA substrates are similar to that of NSUN2’s tRNA substrates. Type II m^5^C sites are adjacent to a downstream UCCA motif and predicted to locate in loops of stem-loop structures. However, the biogenesis of Type II m^5^C sites, as well as the mechanisms responsible for their selective methylation, remains elusive.

## Results

### Computational inference of a Type II m^5^C site-specific methyltransferase

To identify the potential methyltransferase responsible for Type II m^5^C site methylation, we considered deducing the candidate based on the correlation between Type II m^5^C site number and gene expression level. We first confirmed the UCCA motif as a robust signature of Type II m^5^C sites with the low false-assignment rate of Type I m^5^C sites (**Note S1** and **Figure S1A-B**) and then used it to define and characterize Type II m^5^C sites across human, mouse, and fly samples profiled with mRNA BS-seq (**Table S1**). The number of Type II m^5^C sites, as well as their proportions (the number of Type II m^5^C sites divided by the total number of m^5^C sites in a sample), varied across samples, ranging from 5 to 169 sites and from 1% to 40% of the total m^5^C sites (**Figure 1A** and **Table S2**). Additionally, the distributions of the methylation levels of Type II m^5^C sites also differed among samples (**Figure S1C**). Interestingly, both conserved and lineage-specific methylation profiles of Type II m^5^C sites were observed. For example, Type II m^5^C sites were highly enriched in testis in all three species studied, and enriched only in human liver but not in mouse liver. Moreover, Type II and Type I m^5^C sites had an overall similar distribution of genic locations (**Figure 1B**), although compared with Type I m^5^C sites, a slightly lower proportion of Type II m^5^C sites was found in 5’UTR regions in human and mouse (**Figure 1C**).

**Figure 1.**
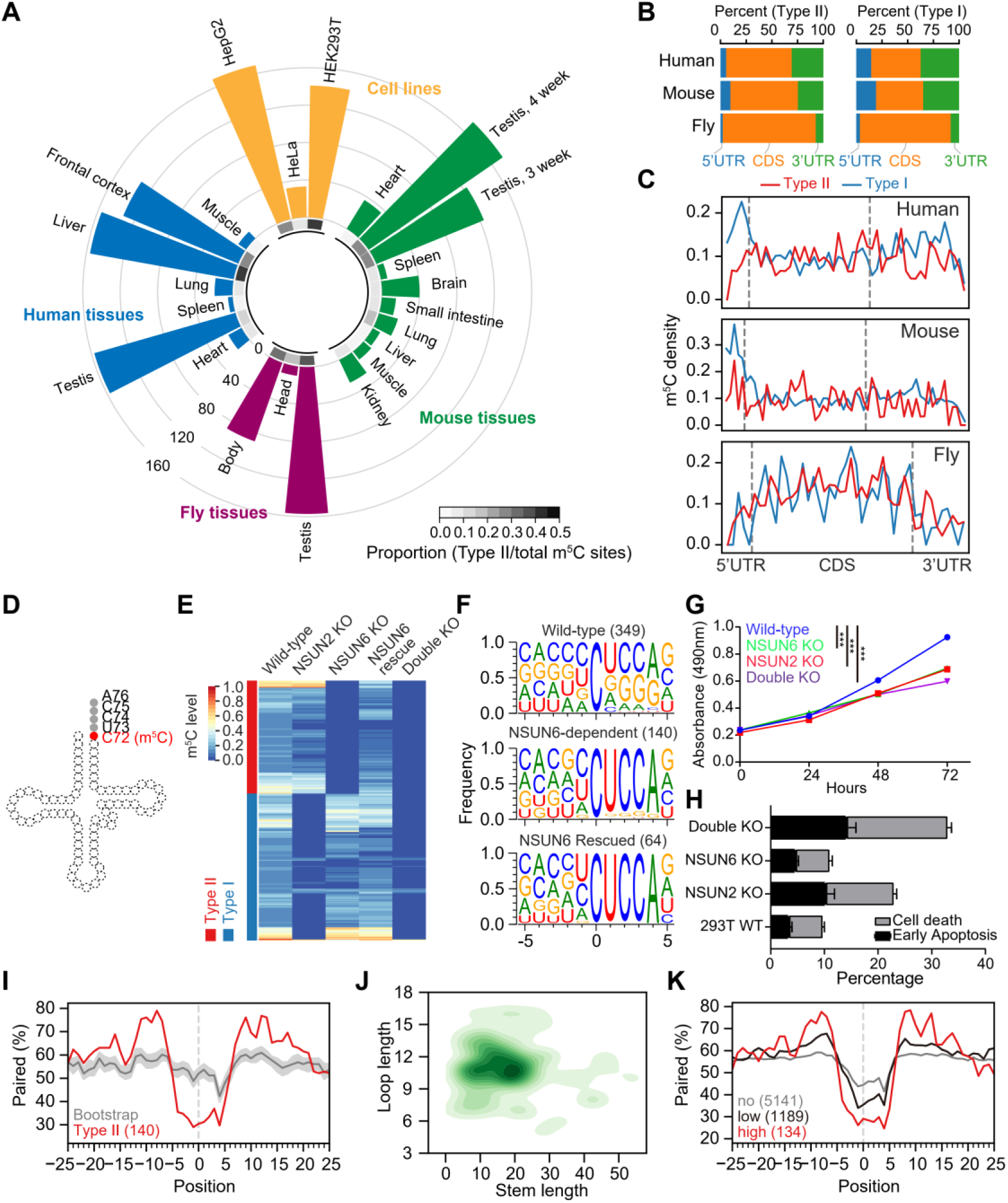
Identification of NSUN6 as an mRNA m^5^C methyltransferase. **(A)** Circular barplot showing the proportion and number of Type II m^5^C sites in cell lines and tissues. Inner circle, the proportion of Type II m^5^C sites in each sample; outer circle, the number of Type II m^5^C sites in each sample. The proportion means the number of Type II m^5^C sites divided by the total number of m^5^C sites in a sample. 7 human and 10 mouse mRNA BS-seq samples from our previous study [17] and 3 wild-type human cell lines and 3 fly tissues mRNA BS-seq samples generated from this study (**Table S2**) were analyzed. **(B)** The genic locations of Type II and Type I m^5^C sites in human, mouse and fly. **(C)** The distributions of Type II and Type I m^5^C sites across the transcripts. The details of the density calculation are described in **Methods**. **(D)** Illustration of the tRNA m^5^C site methylated by NSUN6. **(E)** Heatmap showing the m^5^C methylation levels of sites measured from wild-type, NSUN2 knockout, NSUN6 knockout, NSUN6 rescue, and NSUN6/NSUN2 double knockout HEK293T cells. Sites covered by at least 20 reads in all samples and with methylation level ≥ 0.1 in wild-type cells are shown. **(F)** The sequence context flanking different groups of sites. The number of sites used for analysis in each group is given in parenthesis. NSUN6-dependent, sites that are covered in both wild-type and NSUN6 knockout cells and not methylated (< 0.05) in knockout cells; NSUN6 rescued, sites that are covered in both NSUN6 knockout and rescue cells, not methylated (< 0.05) in knockout cells, and methylated (≥ 0.05) in rescue cells. **(G)** The MTS assay was used to quantify the viable cells in different time points. Data are presented as Mean ± SEM (n = 4). The p values were determined using the Student’s t-test by comparing each of the mutant samples with the wild-type samples at 72 hours. ***, p<0.001. **(H)** Propidium iodide (PI) and Annexin V were used to determine the death or early apoptosis of the cells. The percentage of viability was measured at 48 hours. Data are presented as Mean ± SEM (n = 3). **(I)** Metaprofiles of the secondary structure of NSUN6-dependent m^5^C sites and flanking regions. Position 0 represents the m^5^C sites. Each negative or positive value indicates the distance between an upstream or downstream position and the m^5^C site. The upstream and downstream 50 nt sequences of the m^5^C sites were extracted from the transcriptome and folded with the RNAfold tool, and the 25 nt flanking regions are shown. The same set of NSUN6-dependent sites as in **F** were used for analysis. The same number of Cs with a UCCA motif on m^5^C-containing genes were sampled 100 times (see **Methods**), and the median and the quartiles estimated by bootstrap are shown in gray. **(J)** 2D KDE plot showing the distribution of the stem length and loop length of predicted secondary structures of NSUN6-dependent sites and flanking regions. The same set of NSUN6-dependent sites as in **F** were used for analysis. **(K)** Metaprofiles of the secondary structure of the NSUN6 miCLIP binding sites and flanking regions. Binding sites with a CUCCA sequence were divided into three groups for analysis: no methylation, <1%; low-methylation, 1%-10%; and high-methylation, ≥10%.

Next, we examined the correlation between the expression levels of each of 125 known methyltransferases and the number of Type II m^5^C sites across 9 human samples. Among them, we identified NSUN6, which was originally identified as a tRNA methyltransferase, as the one with the highest correlation coefficient (**Figure S1D, Table S3**). A strong positive correlation was also observed in mouse samples (**Figure S1E**). Interestingly, NSUN6 is known to specifically target the C72 position at the 3’end of tRNA^Thr^ and tRNA^Cys^ with a UCCA tail in humans (**Figure 1D**) [22, 23], in line with the enriched UCCA motif of Type II m^5^C sites. Taken together, these results enabled us to predict that NSUN6 is a Type II m^5^C site-specific methyltransferase.

### Experimental verification of NSUN6 as the methyltransferase responsible for Type II m^5^C sites

To experimentally verify our prediction, we generated NSUN6 knockout human cells (**Figure S2A-B, Table S4**). Human HEK293T cells were selected because of the high proportion of Type II m^5^C sites observed. We found that 49.8% (140/281) of m^5^C sites identified in the wild-type sample were not methylated in the knockout sample (**Figure 1E, Note S2**), and the methylation of these sites was partially rescued by transiently expressing NSUN6 in the knockout cells (**Figure 1E**). A UCCA motif was found to be adjacent to most (123/140) NSUN6-dependent sites (**Figure 1F**), and the loss of methylation on sites with a UCCA motif was further confirmed using NSUN6 knockout HeLa cells (**Figure S2C**). Notably, compared with HEK293T cells, HeLa cells had a lower NSUN6 expression level (**Figure S2D**). Only 30 sites were identified as NSUN6-dependent Type II m^5^C sites in HeLa cells; among them, 27 sites were overlapped with Type II m^5^C sites in HEK293T cells. When both NSUN6 and NSUN2 were knocked out in HEK293T cells (**Figure S2A-B**), nearly all mRNA m^5^C was eliminated (**Figure 1E**). These observations indicate that NSUN6 is an mRNA methyltransferase and, together with NSUN2, determines most mRNA m^5^C.

Characterization of NSUN6 knockout HEK293T cells revealed high viability but reduced proliferation (**Figure 1G-H** and **Figure S2E**). Interestingly, no proliferation defect was observed for NSUN6 knockout HeLa cells (**Figure S2F**), which may be because NSUN6 contributed little to mRNA m^5^C in HeLa cells (**Figure 1A** and **Figure S2C**). It should be noted that we cannot exclude the possibility that the proliferation defects may be due to the lack of tRNA methylation. As a comparison, we examined NSUN2 knockout HEK293T cells and revealed reduced cell viability (**Figure 1G-H**), consistent with the previous studies [24]. Finally, we found that NSUN6/NSUN2 double knockout HEK293T cells exhibited a more severe cell proliferation defect as compared with the single knockout cells (**Figure 1G-H**). These findings highlight the possible physiological importance of NSUN6.

Next, we examined the structural feature of NSUN6-dependent sites. Compared with background cytosines (see **Methods**), NSUN6-dependent sites had a lower percentage of base-pairing in the ∼5 nt flanking regions and a higher percentage of base-pairing in the 10-15 nt flanking regions (**Figure 1I-J**). We then utilized miCLIP data to confirm the NSUN6-mediated methylation of Type II m^5^C sites [7, 25]. As expected, frequent reverse transcription stalling was observed at the NSUN6-dependent but not the NSUN6-independent sites in NSUN6 miCLIP data, and the opposite pattern was observed in NSUN2 miCLIP data (**Figure S2G**). Interestingly, about 59% of the NSUN6 binding sites were adjacent to a downstream UCCA motif. Moreover, the methylation status of these UCCA motif sites was positively correlated with the strength of the predicted stem-loop structure (**Figure 1K**). These results confirmed the direct binding and sequence context requirement for NSUN6-mediated mRNA methylation.

As NSUN6 is found in both vertebrates and invertebrates, we asked whether NSUN6 also methylates Type II m^5^C sites in invertebrates. NSUN6 knockout flies were generated and verified by RT-PCR (**Figure S3A-B)**. We first confirmed that fly NSUN6 also methylated the 3’end of tRNA^Thr^ and tRNA^Cys^ (**Figure S3C**). Next, we examined the mRNA m^5^C methylation. We found that a group of mRNA m^5^C sites, whose sequence and structural features were similar to that of Type II m^5^C sites in mammals, lost their methylation in NSUN6 knockout flies (**Figure S3D-E**), although their motif had a stronger position -1 C preference and a wider base tolerance at position +3. These results are consistent with the predicted conserved NSUN6-RNA complex structure between vertebrates and invertebrates (**Figure S3F**), suggesting NSUN6 as a conserved mRNA m^5^C writer.

### Validation of the sequence and structural requirement for NSUN6-mediated mRNA methylation

To examine the impact of the sequence and structural features identified in our computational model for achieving methylation, we employed a high-throughput mutagenesis assay. One Type II m^5^C site that is highly methylated in HEK293T cells was selected for the assay (**Figure 2A**). On the basis of this target, we designed sequence variants to systematically impact the sequence motif and secondary structure surrounding the m^5^C site (**Table S5**). Each of these sequences further comprised a barcode and common adapters on both ends. The sequences were synthesized as DNA oligo pools, cloned as 3’UTR elements downstream of a luciferase gene, and transfected into wild-type HEK293T cells. Their methylation levels were determined by targeted BS-seq.

**Figure 2.**
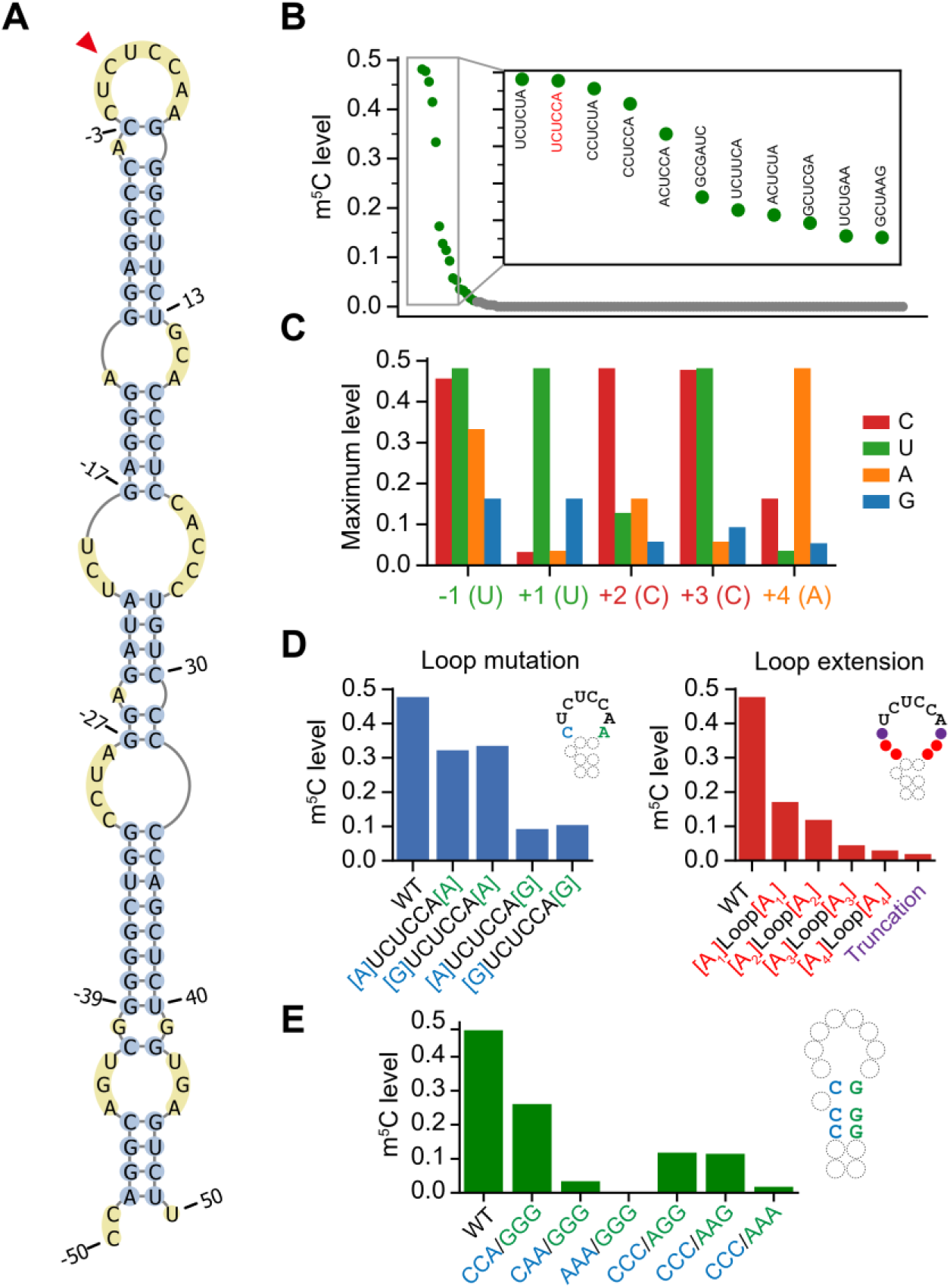
Validation of the sequence and structural requirement for NSUN6-mediated mRNA methylation. **(A)** The predicted secondary structure of an NSUN6-dependent m^5^C site (chr15:91425541) and flanking region. The m^5^C site is indicated by the red arrowhead. **(B)** The NSUN6 core motif (Nm^5^CUCCA) was systematically mutated and the ranked methylation levels of each variant are shown. The original motif of the substrate is colored in red. Y-axis represents the methylation levels of the substrates with the indicated motif sequences. One targeted BS-seq library was constructed. Only motifs with at least 20 reads covered are shown (**Table S2**). **(C)** Barplot showing the maximum methylation levels of the substrates with mutations in the core motif. Y-axis represents the maximum methylation levels of each of the 4 bases in each position. The maximum level was represented using the methylation level of the substrate with the highest methylation level with the indicated base. **(D)** The methylation levels of the substrates with mutation, deletion or extension of the loop region. Truncation, bases at positions -2 and +5 were deleted from the original loop. Y-axis represents the methylation levels of the substrates with the indicated sequences. The number of reads covered for each sequence is shown in **Table S2**. **(E)** The methylation levels of the substrates of which the base-pairing of the stem regions were disrupted by replacing C/G by As. Y-axis represents the methylation levels of the substrates with the indicated sequences. The number of reads covered for each sequence is shown in **Table S2**.

We began by measuring the extent to which each individual nucleotide surrounding the m^5^C site was required for methylation. Overall, except for the original sequence, only several other sequence variants were highly methylated (**Figure 2B**). More specifically, we found that 1) position -1 preferred non-G bases, 2) position +3 favored C and U, 3) original base composition at positions +1, +2, and +4 led to better methylation (**Figure 2C**). We next examined the impact of the more distal flanking bases. Reduced methylation levels were observed when positions -2 or/and +5, which locate in the loop, were mutated, and a more reduced level was observed when position +5 base was mutated (**Figure 2D**). Other efforts were made in modifying the loop and the stem. When positions -2 and +5 were deleted, the substrate completely lost its methylation (**Figure 2D**); when the loop was extended by adenosines, methylation levels were reduced along with the increase of adenosine number (**Figure 2D**); and when more than one base pairing was disrupted, nearly no methylation was observed (**Figure 2E**). These results together suggest both the sequence motif and the stem-loop structure are important for the methylation process.

### Characterizing the NSUN6-mRNA interaction

To further reveal the mechanisms responsible for the sequence- and structure-selective recognition and methylation of mRNA substrates, we sought to use Rosetta comparative modelling [26], reporter assay and *RNP-denovo* [27] to model NSUN6-RNA complex structures in multiple species and identify residues responsible for NSUN6-RNA interaction.

Based on the known human NSUN6-tRNA co-crystallization data (**Figure 3A**) [28], we first modelled the NSUN6-tRNA structures in mouse and fly and examined the residues that were predicted to interact with the core m^5^C motif, particularly positions -1 and +3 of m^5^C sites, which differ between vertebrates (human and mouse) and invertebrates (fly) (**Figure S4A-B**). We found that, compared with human and mouse NSUN6, fly NSUN6 had a more negatively charged methylation pocket (**Figure S4C-D**), which might lead to the stronger C base preference of position -1 in fly. A previous study showed that position +3 (tRNA C75) interacted with the methylation pocket residues, including Tyr131, Lys192, Gly193, and Asp209 [28]. In human and mouse, U was less favored at position +3 because the reduced hydrogen bond might form between U and NSUN6, while A or G at position +3 was not allowed due to steric hindrance (**Figure S4E-F**). In fly, all four bases might be allowed at position +3 because Asp209 and Lys192 were replaced by Gly and Ser, which resulted in a larger space for position +3 bases (**Figure S4E-F**); C remained the most favorable base because it might have more interactions with the residues (**Figure S4F**).

**Figure 3.**
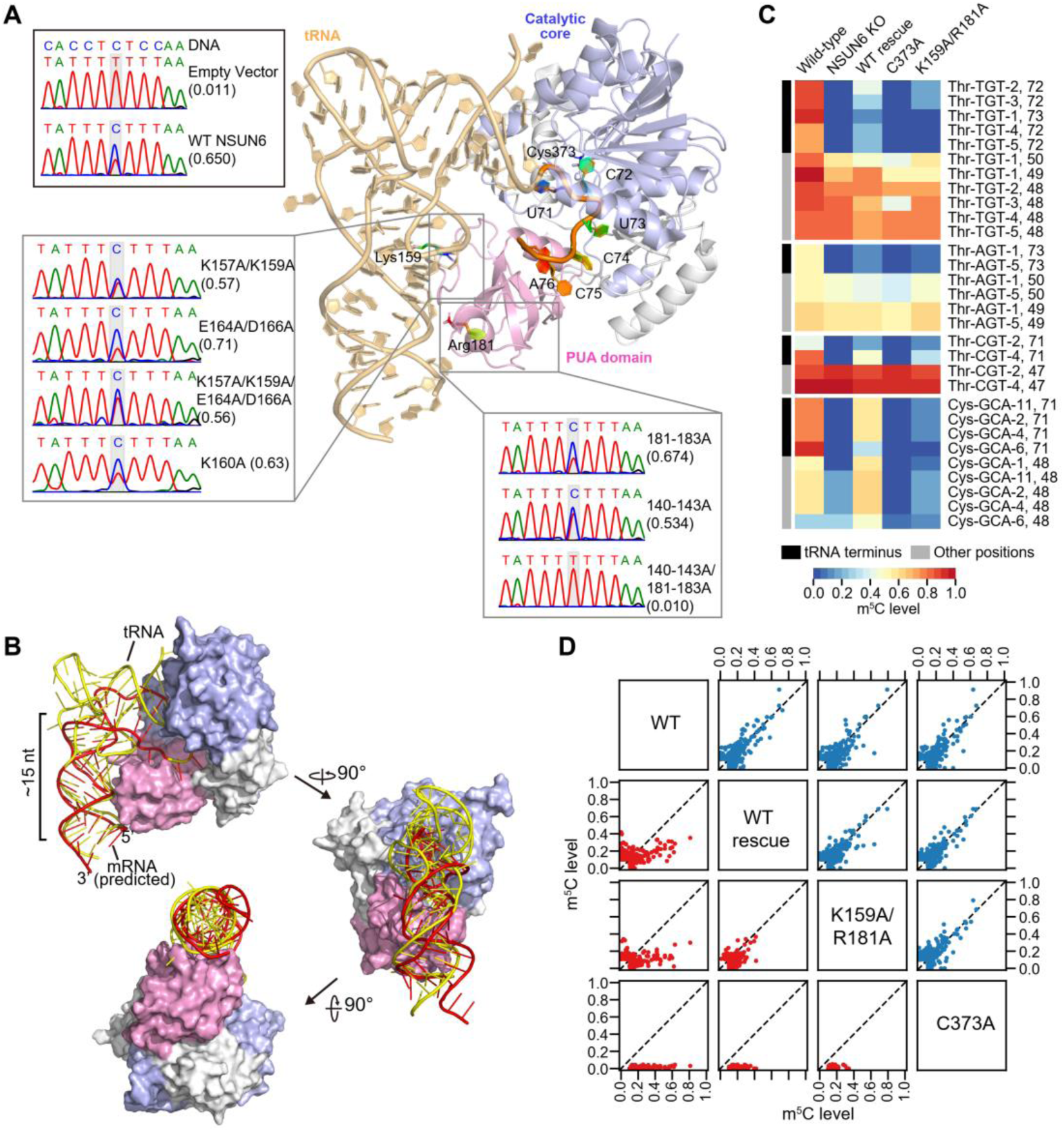
Identification of an NSUN6 variant that largely loses tRNA methylation but retains mRNA methylation ability. **(A)** The overall structure of the human NSUN6-tRNA complex (PDB: 5wws) [28]. The Um^5^CUCCA fragment, MTase domain, PUA domain, and key residues are highlighted. The Sanger sequencing traces for the PCR amplicons of the methylation reporter targeted by different NSUN6 variants are also shown. **(B)** A proposed NSUN6-mRNA structural model in a tRNA-like mode, along with NSUN6-tRNA co-crystallization structure (PDB: 5wws). tRNA is colored in yellow and mRNA is colored in red. **(C)** Heatmap showing the methylation profiles of isodecoders of tRNA^Thr^ and tRNA^Cys^ in the wild-type cells, NSUN6 knockout cells, and NSUN6 knockout cells rescued by wild-type NSUN6 or NSUN6 variants. Sites covered by at least 10 reads in all samples and with a methylation level of ≥ 0.1 in the wild-type cells are shown. **(D)** Pairwise comparison of mRNA methylation levels between the wild-type cells and NSUN6 knockout cells rescued by wild-type NSUN6 or NSUN6 variants. Type I and Type II m^5^C sites are shown in the lower panel (red) and upper panel (blue), respectively. For each pair, sites covered by at least 20 reads in both samples and with a methylation level of ≥ 0.1 in at least one sample are shown.

Next, we examined the interaction between NSUN6 and mRNA stem-loop structure. Given that the PUA domain is responsible for D-stem recognition in tRNA methylation [28] and most mRNA substrates have a stem-loop structure, we predicted that NSUN6 would also interact with mRNA substrates via the PUA domain. Based on the NSUN6-RNA structural model, we performed mutagenesis that might lead to significant shape/electronic changes on the PUA domain (**Figure S4G-H**) and used a Type II m^5^C site reporter (see **Method**) to examine the methylation activity of these NSUN6 variants. We found that 140-143A/181-183A variant, which severely breaks the positively charged distal end surface of the PUA domain, led to a loss of methylation (**Figure 3A**), while other variants retained mRNA methylation activity (**Figure 3A**). These results suggest that, unlike NSUN6-mediated tRNA methylation that is sensitive to single residue mutation [28], mRNA m^5^C is PUA-dependent but less sensitive to specific residues.

Last, we constructed the NSUN6-mRNA complex structure by folding and docking mRNA stem to the NSUN6 protein surface using *RNP-denovo* (**Methods**) and attempted to reveal the possible difference between NSUN6-tRNA and NSUN6-mRNA interactions. We found that one of the best-scored models fitted our current understanding of NSUN6-mRNA recognition: mRNA in a tRNA-like conformation (**Figure 3B**). In this model, the mRNA strand twisted as the tRNA anticodon stem, forming a ∼15 nt stem whose axis overlapped with that of anticodon stem. This model explained why mutations around residues 157-163 retained mRNA methylation activity: although these residues (e.g. Lys159) might be essential for tRNA methylation, they were away from the mRNA strand in the NSUN6-mRNA model.

### Identification of an NSUN6 variant that largely loses tRNA methylation but retains mRNA methylation ability

The results from the reporter assay and NSUN6-mRNA structure modelling suggest that tRNA and mRNA substrates may interact with different residues of NSUN6, thus tRNA methylation and mRNA methylation may be separated by modifying residues in NSUN6. As proof of concept, we selected the known tRNA methylation null mutant K159A/R181A [28] for a test. Given that residues 181-183 do not affect mRNA methylation, we predicted that this mutant may not affect mRNA methylation.

To verify our prediction, we transiently expressed the K159A/R181A mutant in NSUN6 knockout HEK293T cells, using the catalytically inactive C373A mutant and wild-type NSUN6 as the negative and positive controls, respectively, and then conducted mRNA and tRNA m^5^C profiling. For tRNA, as expected, C72 of tRNA^Thr- TGT^, tRNA^Thr-CGT^, tRNA^Thr-AGT,^ and tRNA^Cys-GCA^ completely lost their methylation in both the knockout cells and the C373A mutant rescued cells (**Figure 3C, Figure S5A**, and **Note S3**). Compared with the cells rescued by wild-type NSUN6, very low-levels of methylation were found in the K159A/R181A mutant (**Figure 3C** and **Note S3**). For mRNA, Type II m^5^C sites lost all methylation in both the knockout cells and the C373A mutant rescued cells, but similar Type II m^5^C site methylation profiles were observed between the K159A/R181A rescued cells and the wild-type NSUN6 rescued cells (**Figure 3D**). Notably, both the wild-type NSUN6 and the K159A/R181A mutant only partially rescued the methylation levels of their mRNA targets. Western blot analysis revealed that the exogenous NSUN6 proteins had even a higher level than the endogenous one (**Figure S5B**), suggesting that their activities might be affected by unknown factors. These findings suggest that tRNA methylation and mRNA methylation may be distinguished via use of an edited NSUN6 protein.

### NSUN6 and NSUN2 methylate mRNAs in different cellular compartments

Next, we sought to assess within which subcellular compartments Type II and Type I m^5^C sites are localized. We purified nuclear and cytoplasmic-enriched subcellular fractions in HEK293T cells and performed mRNA BS-seq on these fractions. The purity of the nuclear and cytoplasmic fractions was confirmed by quantitative PCR (qPCR) and western blot (**Figure S6A-C**). We found that m^5^C levels of a substantial fraction of Type II m^5^C sites were higher in the cytoplasm than that in the nucleus, while most Type I m^5^C sites had similar m^5^C levels between the two subcompartments (**Figure 4A**). Moreover, when examining the m^5^C sites called from nuclear RNA, only Type I m^5^C sites were observed in introns (**Figure 4B**). At the protein level, in HEK293T cells, NSUN2 was mainly located in the cytoplasm; in HeLa and HepG2 cells, NSUN2 was located in both the nucleus and cytoplasm (**Figure S6C**). NSUN6 was mainly located in the cytoplasm (**Figure S6C**), consistent with the known subcellular location of NSUN6 in the Golgi apparatus [22]. In line with this, NSUN6 miCLIP data revealed that more than 89.8% of the NSUN6 binding sites were located in exonic regions. These analyses together suggest that NSUN6 may methylate mRNAs in the *cis*-Golgi network, while NSUN2 may methylate mRNAs mainly but not exclusively in the nuclear subcompartment.

**Figure 4.**
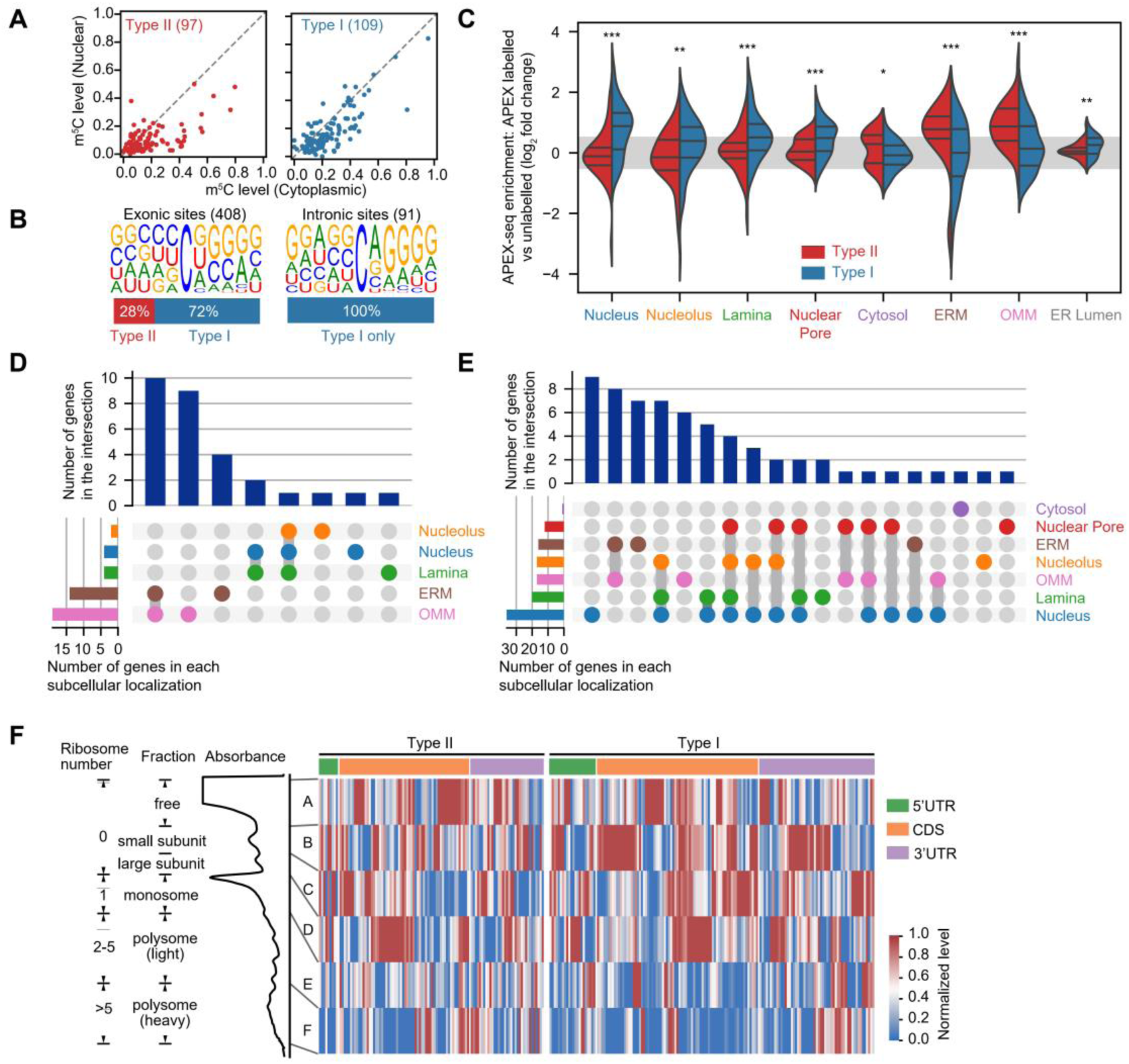
Characterizing the deposition and the molecular readout of Type II and Type I m^5^C sites. **(A)** Comparison of methylation levels of m^5^C sites measured in nuclear and cytoplasmic poly(A)+ RNAs. Sites covered by at least 20 reads in both samples are shown. **(B)** Motif analysis for m^5^C sites called from nuclear poly(A)+ RNAs. Exonic sites and intronic sites were analyzed separately. The percentages of Type I and Type II m^5^C sites in exonic and intronic sites are also shown. **(C)** Violin plot showing the enrichment of genes containing Type I or Type II m^5^C sites in different subcellular localizations. HEK293T APEX-seq data were used for analysis [29]. Y-axis represents the APEX-seq enrichment, which means log2fold change over the corresponding negative controls. Inner bars, median and two quartiles. The significant difference between genes containing Type II and Type I m^5^C sites was determined using a two-sided Mann−Whitney U test. *, p < 0.05; **, p < 0.01; ***, p < 0.001. (**D-E**) UpSet plot showing the overlapping number of genes with Type II (**D**) or Type I (**E**) m^5^C sites in different subcellular localizations. UpSet plots the intersections of a set as a matrix. Each row corresponds to a set (subcellular localization). Cells are either empty (light-gray), indicating that this set is not part of that intersection, or filled, showing that the set is participating in the intersection. Therefore, the barplot on the top shows the number of genes in the intersection; the barplot on the bottom shows the number of genes in each subcellular localization. (**F**) Methylation levels of sites across the polysomal fractions. Absorbance trace (260 nm) across the gradients processed for RNA BS-seq is shown above the heatmap. Each row represents a Type I or Type II m^5^C site. 154 and 106 Type I and Type II sites with at least 20 reads in at least 5 fractions are shown. The data were Min-Max normalized. In brief, for each site, the original methylation levels were normalized to interval [0,1], according to the following equation: 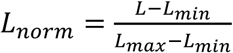, where L is the original value, L_min_ is the minimal value of the original value, L_max_ is the maximal value of the original value and L_norm_ is the normalized value.

### The potential function and molecular readout of Type II m^5^C methylation

The APEX-seq, a method for direct proximity labeling of RNA using the peroxidase enzyme APEX2, has been recently developed and enabled the study of the enrichment of mRNA species in different subcellular localizations [29]. We thus utilized the published APEX-seq data in nine distinct subcellular localizations of HEK293T cells to examine the subcellular localizations of m^5^C-containing transcripts. We found that transcripts containing Type I m^5^C sites were mainly enriched in the nuclear fractions (nucleus, nucleolus, lamina, and nuclear pore) (**Figure 4C-E**). Transcripts containing Type II m^5^C sites were mostly enriched near the endoplasmic reticulum membrane (ERM) and outer mitochondrial membrane (OMM) but not in the endoplasmic reticulum lumen (**Figure 4C-E**). This finding suggests that NSUN6-mediated methylation might be involved in mitochondrial functions.

As a negative correlation between translational efficiency and the presence of mRNA m^5^C was reported previously [17], we asked whether mRNA m^5^C plays a role in translational control. We fractionated the lysate of wild-type HEK293T cells to isolate mRNAs associated with different numbers of ribosomes, and performed mRNA BS-seq in each fraction. A comparison of each fraction revealed that the highest methylation levels of m^5^C sites were more frequently found in mRNAs not bound by the ribosome, and the lowest methylation levels of them were more frequently found in mRNAs bound by multiple ribosomes (**Figure 4F** and **Figure S7A**). This result confirmed a negative correlation between m^5^C methylation and translation efficiency *in vivo*. To further measure the impact of m^5^C methylation on translational control, we focused on Type II m^5^C sites and selected 3 sites (2 in CDS and 1 in 3’UTR) with high-methylation in HEK293T cells for a dual-luciferase reporter assay. We cloned individual m^5^C sites and flanking regions into the reporter gene and generated C-to-T point-mutation as a control (**Table S6, Methods**). We first confirmed that the wild-type plasmids were methylated when expressed in HEK293T cells (**Figure S7B**). Next, each of the wild-type or mutated plasmids was transfected into the wild-type or NSUN6 knockout HEK293T cells. We found that, in all cases, methylated transcripts had a slightly decreased luciferase level (**Figure S7C-D, Table S7**). We also examined the protein expression of genes with selected m^5^C sites in the wild-type and NSUN6 knockout cells. We found that 1 gene tended to have a decreased protein expression in the wild-type cells, although it had a slightly increased RNA level in the wild-type cells (**Figure S7E-F**). These results together suggested a weak negative correlation between mRNA m^5^C and translational efficiency, and the underlying mechanism and biological significance need to be investigated in future studies.

## Discussion

In this study, we identified NSUN6 as an mRNA methyltransferase. We found that NSUN6 specifically methylated a group of sites in a sequence- and structure-selective manner. Notably, NSUN6 showed a tissue-specific expression pattern, with higher expression levels in testis, pituitary, ovary, liver and thyroid (https://www.proteinatlas.org/ENSG00000241058-NSUN6/tissue). Consistently, more Type II m^5^C sites were observed in tissues with high-level NSUN6 expression (**Figure 1A**). Our findings pave the path toward mechanistic dissection of the physiological importance of NSUN6, such as its roles in cell proliferation and tumor progression [30, 31].

NSUN6 is a methyltransferase known to methylate C72 of tRNA^Thr^ and tRNA^Cys^. Compared with its tRNA methylation sites, most of NSUN6’s mRNA sites had much lower methylation levels. For example, of the 503 Type II sites found in at least one of the 10 human tissue and cell samples (**Figure 1A**), only 61 sites had a methylation level > 0.4 in one or more samples. Interestingly, it is known that archaeal NSUN6 has a much wider range of tRNA substrates [32], but both human and fly NSUN6 proteins only methylated 4 tRNAs at C72. Moreover, miCLIP data revealed that about 30% of human NSUN6 miCLIP reads were mapped to mRNAs. These data suggest that NSUN6 in higher animals may be evolving to bind fewer tRNAs but more mRNAs, although many of the mRNA substrates are still lowly methylated.

One of the most challenging problems in the RNA modification field is that most RNA modification writer proteins have multiple types of RNA substrates, thus it is difficult to study the specific functions of one type of RNA substrates. Here using NSUN6 as a model, we demonstrated that, based on protein-RNA structure analysis, we were able to identify an NSUN6 variant that largely loses tRNA methylation but retains mRNA methylation ability. We anticipate that our approach is applicable to other writer proteins as well, such as NSUN2 (m^5^C), METTL16 (m^6^A), TRMT6/TRMT61A (m^1^A) and PUS proteins (pseudouridine). Similar to NSUN6, although their mRNA substrates mimic the features of tRNA or snRNA [4], they are likely to have more flexible structural or sequence features and their methylation may require a less stringent protein-RNA interaction. Therefore, it is possible to generate variants that lose tRNA or snRNA modification but retain mRNA modification ability. And these variants may enable the specific investigations of the functions of their mRNA substrates.

## Acknowledgments

We thank Dr. Chung-I Wu for sharing the equipment and space for fly experiments, Dr. Xiaohang Yang for sharing the *D*.*mel* CRISPR/Cas9 tool stains, Dr. Michaela Frye, Dr. Holger Prokisch and Dr. Michal Minczuk for the generation of NSUN6 and NSUN2 miCLIP data (GSE66011 and GSE44386), and SYSU Ecology and Evolutionary Biology Sequencing Core Facility for the sequencing service. This study was supported by grants from Guangdong Major Science and Technology Projects (2017B020226002 to R.Z.), National Key R&D Program of China (2018YFC1003100 to R.Z.), Guangdong Innovative and Entrepreneurial Research Team Program (2016ZT06S638 to R.Z.), National Natural Science Foundation of China (31571341 and 91631108 to R.Z.; 41031230 to T.H.), Science Foundation for Youth Scholars of Ministry of Education of China (31611190 to T.H.) and China Postdoctoral Science Foundation (41091272 to T.H.).

## Author contributions

T.H., J.H.L., Y.S.Z., T.X.Z, X.N.Z, and W.Y.C. conducted the experiments. J.H.L performed the bioinformatics analysis. J.H.L, T.H., and R.Z. wrote the manuscript with input from all authors.

## Declarations of interests

The authors declare no competing financial interests.

## METHODS

### Cell culture

HEK293T, HeLa and HepG2 cell lines were purchased from Cell Bank, Type Culture Collection, Chinese Academy of Sciences (CBTCCCAS). All cell lines have been identity verified using short tandem repeat (STR) analysis, which involves the simultaneous amplification of 17 STR markers plus amelogenin to confirm the identity of the cells, by CBTCCCAS. Cell lines were maintained in DMEM (Gibco) supplemented with 10% FBS (HyClone). All cell lines have been checked for mycoplasma contamination by CBTCCCAS and are routinely tested for mycoplasma by PCR detection of conditioned medium.

### Fly husbandry

Fly stocks were kept at 25 °C on a 12 hours light and 12 hours dark cycle and fed cornmeal food.

### NSUN6 plasmid construction

To construct NSUN6 expression plasmid, cDNA from HEK293T cells was reverse transcribed using HiScript II Q RT SuperMix (Vazyme), and the full-length NSUN6 CDS fragment was amplified. The NSUN6 fragment was inserted into the AvrII and AgeI sites of the pCDH-3xFLAG vector to generate pCDH-3xFLAG-NSUN6 plasmid. NSUN6 mutagenesis was performed using Mut Express II Fast Mutagenesis Kit V2 (Vazyme) or CloneExpress Ultra One Step Cloning Kit (Vazyme).

### Knockout cell generation and rescue experiment

NSUN2 knockout, NSUN6 knockout, and NSUN2/NSUN6 double knockout cells were generated via CRISPR/Cas9-induced mutagenesis. In brief, sgRNA sequences were designed using CRISPR-ERA (http://CRISPR-ERA.stanford.edu) (**Table S4**). The sgRNA template oligonucleotides were synthesized and cloned into lentiCRISPR v2 plasmid (Addgene#52961). The plasmid was transfected into the cells using Lipofectamine 3000 (Thermo Fisher Scientific) following the manufacturer’s instruction. Transfected cells were selected using puromycin (1μg /ml). Mutant clones were selected by Sanger sequencing.

For the rescue experiment, knock out cells were plated in a 6-well plate. 3 μg of NSUN6 plasmid was transfected using Lipofectamine 3000 following the manufacturer’s instruction. 48 hours after the transfection, proteins of the cells were collected to confirm the expression of NSUN6 by western blot.

### Nuclear/cytoplasmic fraction

HEK293T cells were fractionated using the PARIS kit (Ambion) according to the manufacturer’s instruction. The separation of cytoplasmic and nuclear fractions was validated using qPCR and western blot. For qPCR, the cytoplasmic and nuclear RNAs were reverse transcribed using HiScript II Q RT SuperMix (Vazyme), and the qPCR was performed using ChamQ SYBR qPCR Master Mix (Vazyme) with the following primers: the exon 8 of PKM: AGAGCATGATCAAGAAGCCCC (forward) and GAGGATAGTCCCCTTTGGCTG (reverse); the intron 7 of PKM: ACACTCGCATGTTTGTATGGG (forward) and TGTTACGTGCGACAATTCCA (reverse). For western blot, GAPDH and LaminB1 were used as cytosolic and nuclear markers, respectively.

### *Drosophila* genetics and sample collection

#### Drosophila melanogaster

W1118 was used as wild-type control. The mutant allele for NSUN6 was generated using the CRISPR/Cas9-induced mutagenesis system following the previously described procedure [33]. In brief, a sgRNA sequence was designed using flyCRISPR Optimal Target Finder (http://www.flyrnai.org/crispr/) (**Table S4**). The sgRNA template oligonucleotide was synthesized and cloned into the donor vector pUAST-attB. The sgRNA-pUAST-attB plasmid was microinjected into 86Fb (Bloomington 24749) embryos. The microinjection was performed by Core Facility of Drosophila Resource and Technology, SIBCB (CAS). The sgRNA transgenic line was crossed with vas-Cas9, so the progeny expressed both sgRNA and vas-induced Cas9 in the germline. The genotype of F1 was screened by Sanger sequencing.

### Construction of NSUN6 substrate plasmids

For low-throughput experiments, the substrate template oligos were synthesized and individually cloned into the psiCHECK-2 vector using the seamless cloning method with CloneExpress Ultra One Step Cloning Kit (Vazyme). Plasmids were extracted with the endotoxin-free plasmid extraction kit (TIANGEN).

For high-throughput experiments, the substrate template oligos were first annealed and extended to form a ∼150 bp double-stranded insert fragments (**Table S5**). The reactions were performed using NEBuilder HiFi DNA Assembly Master Mix (NEB) by incubation at 50 °C for 15 minutes. The insert fragments were then purified and mixed with the linearized vector in a 7:1 molar ratio. 4 µl assembled products were used for bacteria transformation. Bacteria were shaken in SOC medium in 250 rpm for 60 minutes and then plated on the 14 cm LB agar plates. After incubation for about 14 hours, 2 plates of bacteria (∼20,000 colonies per plate) were harvested. Plasmids were extracted with the endotoxin-free plasmid extraction kit (TIANGEN).

### Analysis of cell viability and apoptosis

For cell viability analysis, we used the CellTiter 96 AQueous One Solution Cell Proliferation Assay kit (Promega) according to the manufacturer’s instruction. In brief, 1 x 10^3^ wild-type or knock out cells were seeded into each 96-well plate. 0, 24, 48 and 72 hours after the seeding, 20 μl One Solution Reagent was added. The cells were incubated at 37 °C for 1 hour, and the absorbance at 490 nm was recorded.

For cell apoptosis analysis, Annexin V -FITC and Propidium iodide (PI) staining were used to determine the early apoptosis and cell death.

### RNA-seq

For each sample, 1 μg total RNA was used for library construction. poly(A)+ RNA was separated from total RNA using Oligo dT Magnetic Beads (Vazyme). RNA was then used for library construction with NEBNext Ultra II Directional RNA Library Prep Kit (NEB). Libraries were sequenced on Illumina Hiseq X10 (Illumina) to produce paired-end 150 bp reads.

### mRNA BS-seq

mRNA BS-seq library construction was performed as we previously described [17]. In brief, total RNA was isolated with TRIzol reagent and Direct-zol RNA MiniPrep kit. poly(A)+ RNA was separated from total RNA using Oligo dT Magnetic Beads (Vazyme). 100 ng - 1 μg of poly(A)+ RNA was converted using the EZ RNA methylation kit (Zymo Research) with a modified high-stringency conversion condition (Sulfonation: 3 cycles, (1) 70 °C, 10 minutes. (2) 64 °C, 45 minutes. Desulfonation: 25 °C for 30 minutes.). The converted RNA was fragmented into 150 - 200 nt fragments by incubation at 94 °C for 8 minutes in fragmentation buffer (NEB). The fragmented RNA was then used for library construction using NEBNext Ultra II Directional RNA Library Prep Kit. SYBR Green I was added to the PCR mixture and the amplification was done with a qPCR instrument so that the amplicons can be monitored at each cycle to avoid unnecessary extra amplification. Libraries were sequenced on Hiseq X10 (Illumina) to produce paired-end 150 bp reads. All libraries are summarized in **Table S1**.

### BS-PCR and targeted BS-seq

Total RNA was extracted with TRIzol reagent and Direct-zol RNA Kit (Zymo Research) 48 hours after the transfection. Total RNA was treated with DNase I, BS converted (Sulfonation: 1 cycle, (1) 70 °C, 10 minutes. (2) 64 °C, 45 minutes. Desulfonation: 25 °C for 30 minutes.), and reverse transcribed with gene-specific primers using HiScript II Q RT SuperMix (Vazyme). Target sequences were amplified using STARmix Taq DNA Polymerase (GenStar) with the following program: 94 °C for 3 minutes; 30 cycles of 94 °C for 30 seconds, 52 °C for 30 seconds and 72 °C for 20 seconds; and 72 °C for 1 minute. For BS-PCR, Sanger sequencing of PCR amplicons was applied to measure the methylation level of the mRNA substrate. For targeted BS-seq or targeted sequencing of plasmid pools, the first round of PCR was performed with 25 cycles using the same program above. Next, the PCR product was purified with DNA Clean Beads (Vazyme) and dissolved in 10 µl water. 1 µl purified product was amplified using sequencing adaptors with the same program above for 5 cycles. Finally, the PCR product was recovered with Zymoclean Gel DNA Recovery Kit (Zymo Research) and sequenced. All primers used for targeted BS-seq are listed in **Table S5**.

### tRNA BS-seq

20 µg total RNA was first loaded into 15% denaturing TBE-Urea PAGE Gels and separated based on the molecular weight. The gel slice corresponding to 70-90 nucleotides was excised from the PAGE gel and the RNA fragments were recovered using ZR small-RNA PAGE Recovery Kit (Zymo Research) following the manufacturer’s protocol. RNA was then converted using the EZ RNA methylation kit (Zymo Research) as we previously described [17]. End-repair was performed with T4 PNK (NEB) to phosphorylate 5’-hydroxyl termini and remove 2’,3’-Cyclic phosphate produced during the bisulfite/desulfonation reaction. The tRNA BS-seq library was generated using VAHTS Small RNA Library Prep Kit for Illumina (Vazyme) according to the manufacturer’s protocol. Libraries were sequenced on Hiseq X10 (Illumina) to produce paired-end 150 bp reads. All libraries are summarized in **Table S1**.

### Polysome RNA BS-seq

Cells were collected and lysed with lysis buffer (10 mM Tris-HCl (pH 7.4), 5 mM MgCl_2_, 100 mM KCl, 1% Triton X-100, 2 mM DTT, 100 μg/ml CHX (CST), 50 U/ml RiboLock RNase Inhibitor (Thermo Fisher Scientific) and cOmplete, EDTA-free Protease Inhibitor Tablets (1 tablet per 10ml, Roche)). 10–50% sucrose gradients were prepared in gradient buffer (20 mM HEPES-KOH, pH7.4, 5 mM MgCl_2_, 100 mM KCl, 2 mM DTT, 100 μg/ml CHX, 50 U/ml RiboLock RNase Inhibitor) using a Gradient Master (Biocomp). 400 μl lysates were loaded on the gradients. Gradients were centrifuged at 36,000 rpm for 2 hours at 4 °C in an SW-40 Ti rotor and then fractionated using the Gradient Master. RNA was equally divided into 22 fractions and these fractions were mixed: fraction 1-4, free RNA, with 0 ribosome; fraction 5-6, RNA with ribosome small subunit and 0 ribosome; fraction 7-8, RNA with ribosome large subunit and 0 ribosome; fraction 9-11, monosome; fraction 12-17, light polysome, with 2-5 ribosomes; fraction 18-22, heavy polysome, with >5 ribosomes. RNA was then isolated using TRIzol reagent. poly(A)+ RNA of each fraction was separated from total RNA using Oligo dT Magnetic Beads (Vazyme). About 10-50 ng mRNA of each fraction was converted using the EZ RNA methylation kit (Zymo Research) with a modified high-stringency conversion condition. The converted RNA was then used for library construction with NEBNext Ultra II Directional RNA Library Prep Kit. Libraries were sequenced on Hiseq X10 (Illumina) to produce paired-end 150 bp reads. All libraries are summarized in **Table S1**.

### Dual-luciferase reporter assay

The wild-type fragments were amplified with Phusion High-Fidelity DNA Polymerase (NEB) using HEK293T cDNA. For each of the sites in CDS, a fragment in the m^5^C-containing gene in-frame (BBS4,90 bp; PLTP, 81 bp) was inserted right behind the start codon of the Firefly luciferase [2]. For the site in 3’UTR, a 160 bp fragment harboring the m^5^C site was inserted into downstream of the Renilla coding region. The C-to-T point mutation was introduced using Mut Express II Fast Mutagenesis Kit V2 (Vazyme). Plasmids were transfected into NSUN6 knockout or wild-type HEK293T cells using Lipofectamine 3000. Renilla and Firefly luminescence was measured 24 hours later using Dual-Glo Luciferase Assay System (Promega) on GloMax -96 Microplate Luminometer (Promega). All primers used to construct the reporter genes and quantify the expression of Renilla and Firefly luciferases are listed in **Table S6**.

### Genome assembly and gene models

Genome, transcriptome and gene annotations of human GRCh37.75, mouse GRCm38.85 and fly BDGP5.78 were downloaded from Ensembl. tRNA isodecoder sequences were downloaded from GtRNAdb [34]. “CCA” tails were added before index build. rRNA sequences were downloaded from SILVA database (release 138) [35].

### mRNA BS-seq data analysis

mRNA m^5^C sites were called as we previously described [17]. We first trimmed adapters, the first 10 bp of the reads, the last 6 bp of the reads, and the low-quality bases using Cutadapt (-e 0.25 -q 25 -trim-n) [36] and Trimmomatic [37]. Then clean reads were mapped to the *in silico* converted genome by HISAT2 (-k 10,–fr,–rna-strandness FR,–no-mixed) [38] to obtain unique alignments. The remaining unmapped and multiple mapped reads were further mapped to the *in silico* converted transcriptome by Bowtie2 (-end-to-end,–fr,–gbar 5,–mp 5, -k 10, -R 2, -D 5) [39]. Alignment results were merged together, and only bases with high quality (Q ≥ 30) were used for the variant calling. Last, the sites were called using a series of filters as previously described [17]. In brief, we inspected all positions with C-to-T mismatches and only took variant positions into consideration if they conformed to our requirements for number, frequency, and quality of bases that vary from the converted reference sequences: (i) each variant is supported by 3 or more variant nucleotides having a base quality score of ≥ 30, mismatch frequency ≥ 0.1 and coverage of C+T ≥ 20; (ii) the variant still satisfies the above criteria after the removal of the overlapped C-reads based on the Gini coefficient determined C-cutoff filter; (iii) the signal ratio of the variant is ≥ 0.9; (iv) the variant is not located at conversion-resistant genes and (v) the p-value calculated using one-sided binomial test based on gene-specific conversion rate is < 0.001. The methylation level (mismatch frequency) is defined as the number of reads with C divided by the number of reads with C or T.

### Targeted BS-seq data analysis

Reads were mapped to the reference sequences (DNA library against the original reference sequences; BS-seq library against the C-to-T converted reference sequences) with Bowtie2 (--norc). Custom scripts were used to extract reads with barcodes that are unique in both DNA and BS-seq libraries. The C/T counts at m^5^C position were extracted and the methylation level is defined as the number of reads with C divided by the number of reads with C or T.

### tRNA BS-seq data analysis

Adapters were first trimmed with Cutadapt (--max-n 1 -m 18 -e 0.25 --trim-n -q 20) [36]. Only reads containing adapters were retained. Clean reads were first C-to-T (read 1) or G-to-A (read2) converted and mapped to C-to-T converted reference sequences containing tRNA, mRNA, and rRNA with Bowtie2 (--end-to-end --no-mixed --norc -k 50 -X 80). Read pairs were ranked by alignment scores (sum of AS and YS tags in the BAM file). Only read pairs with the highest alignment score and uniquely mapped to one tRNA type (multiple alignment on tRNA isodecoders was allowed) were used. Last, original reads were recovered with a custom script and piled up to call m^5^C sites and calculate their methylation levels. This method ensures us to discover the loss of methylation at isodecoder level, although it is imprecise in detecting the increase of methylation due to the possible cross-alignment.

### Motif analysis

The m^5^C sites and flanking regions were extracted from the transcriptome (exonic sites) or the genome (intronic sites). Motif logos were plotted with WebLogo [40].

### m^5^C density calculation

To generate the metagene profile of m^5^C site distribution across transcripts, 5’UTR was fixed to 5 bins and then CDS and 3’UTR were divided into bins based on their relative average lengths to 5’UTR (human CDS, 25 bins; human 3’UTR, 20 bins; mouse CDS, 30 bins; mouse 3’UTR, 30 bins; fly CDS, 30 bins; fly 3’UTR, 15 bins). Last, m^5^C density in each bin was defined as the ratio of m^5^C number to the total C number.

### Background C generation

Background cytosines mean Cs adjacent to a UCCA motif on m^5^C-containing genes. Background sites were randomly selected by the bootstrap method. In brief, Cs with UCCA motif were randomly selected from transcripts with Type II m^5^C sites (e.g. if a transcript had 2 m^5^C, 2 Cs were sampled). Then the sequence flanking those sampled Cs were folded. This step was repeated 100 times to calculate the median and quartile for each of the bases.

### Analysis of miCLIP data

NSUN6 (GSE66011) and NSUN2 (GSE44386) miCLIP data were obtained from GEO [7, 25]. 3’ adapters were trimmed with Cutadapt (-m 18 -q 20 --trim-n). Possible 5’ adapters were also trimmed. Clean reads were first mapped to rRNAs (Silva-release132 and Ensembl rRNA and mt-rRNA) with Bowtie2 (--norc -N 0 -L 20). Unmapped reads were further mapped to tRNAs (GtRNAdb and Ensembl tRNA and mt-tRNA). The remaining unmapped reads were mapped to the reference genome and transcriptome (Ensembl release 75) with HISAT2 and only uniquely mapped reads were used for binding site identification in mRNAs. For NSUN2 data, 9%, 81% and 10% of the reads were mapped to rRNA, tRNA, and mRNA, respectively; for NSUN6 data, 31%, 40% and 29% of the reads were mapped to rRNA, tRNA, and mRNA, respectively.

For mRNA binding site identification, the miCLIP truncation sites (5’ ends of aligned reads) within protein-coding gene exons (2,2810 genes for Ensembl release 75) were assigned to the nearest cytosine at ±2nt regions (priority: 0, -1, +1, -2, +2), as previously described [25]. The priority of assignment is: 0, -1, +1, -2, +2. FDR was calculated as previously described [41] but without crosslinking site extension. To generate a background, truncation sites within a gene were randomized and then assigned to the nearest cytosine. Cytosines with FDR < 0.05 were considered as NSUN6 binding sites. For NSUN6 data, sites shared in all replicates were used for analysis. For NSUN2 data, due to the limited number of mapped reads in mRNAs, sites identified in each replicate were merged for analysis. Notably, since these are overexpression experiments, caution is needed when interpreting the results.

### Comparative modelling of mouse and fly NSUN6

General RosettaCM protocol [26] was used in NSUN6 structure prediction. The protein part of NSUN6-tRNA complex (PDB: 5wws) was used as the template. Mouse (ENSMUSG00000026707) and fly NSUN6 (FBgn0037200) protein sequences were first aligned to human NSUN6 (ENSG00000241058) by MUSCLE [42]. 3- and 9-mer fragments of mouse or fly NSUN6, as well as the predicted secondary structure, were generated by Robetta (http://robetta.bakerlab.org/fragmentsubmit.jsp). Then threading protocol was run with minirosetta. 1,000 structures generated by random seeds were ranked and the best ones were selected.

### RNA structure prediction and illustration

RNA secondary structure was predicted using RNAfold (2.4.12) [43] with default settings. The RNA structure in **Figure 2A** was drawn by PseudoViewer 3 web application [44].

### Protein structure illustration

Human NSUN6-tRNA co-crystallization data by Liu et al.[28] were downloaded from Protein Data Bank (PDB: 5wws). Protein annotation of NSUN6 was referred to Liu et al. Protein alignment and structure visualization were performed with PyMOL (http://www.pymol.org/). The electrostatic surface was predicted with APBS electrostatics [45].

### RNA-seq analysis

Adapters were first trimmed with Cutadapt [36]. The clean reads were mapped to the reference genome by Tophat2 [46] in strand-specific manner (--library-type fr-firststrand). Alignments were then processed to HTSeq-count [47] to obtain read counts of each gene. Finally, differential gene expression analysis was performed with DESeq2 [48].

---

**Figure S1.**
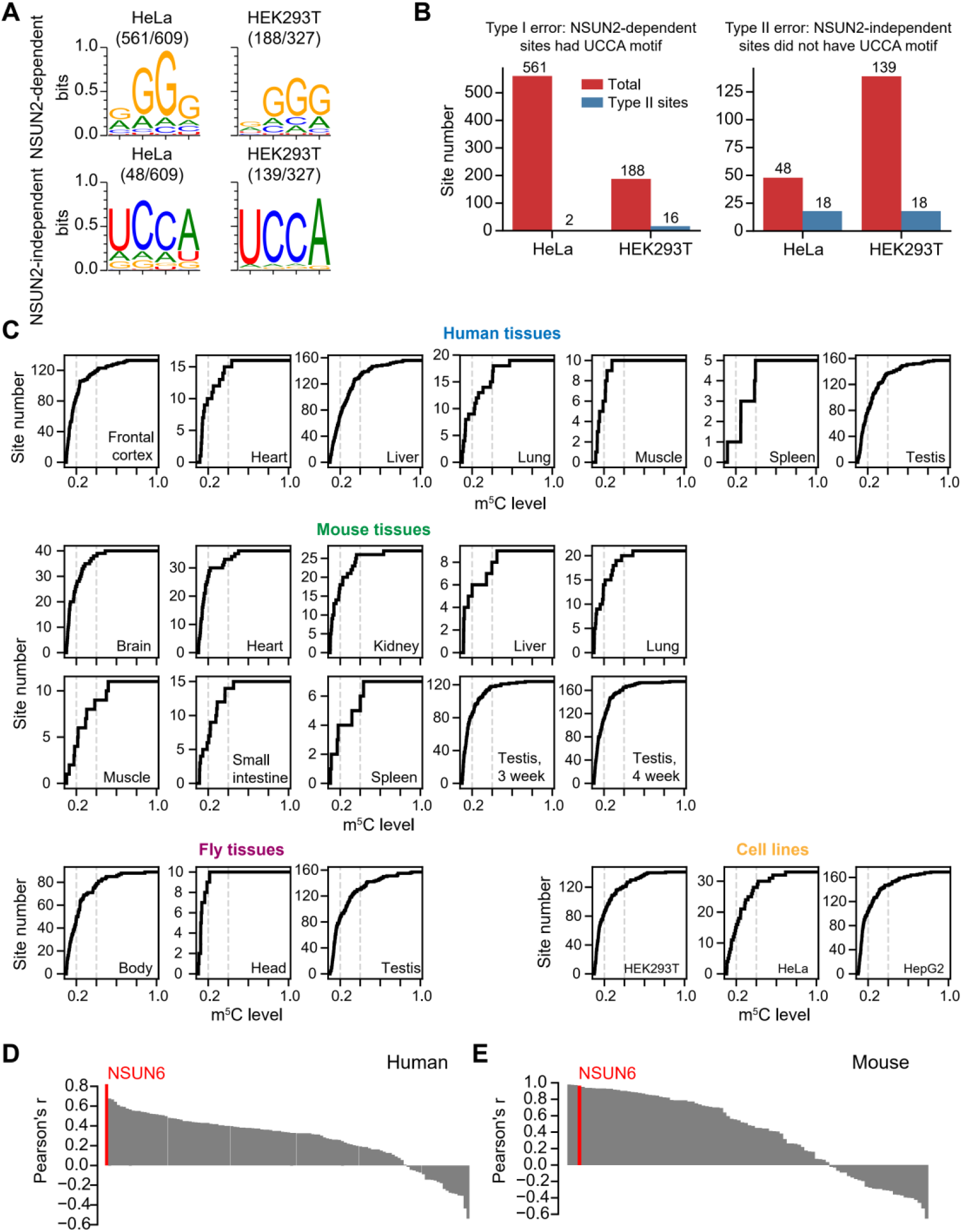
Computational inference of a Type II m^5^C site-specific methyltransferase. **(A)** The sequence motif surrounding NSUN2-dependent and NSUN2-independent sites called from HeLa and HEK293T cells. NSUN2-dependent sites, m^5^C sites identified in wild-type cells but not methylated (m^5^C level < 5%) in NSUN2 knockout cells; NSUN2-independent sites, m^5^C sites methylated in both wild-type and NSUN2 knockout cells. HeLa cell data were from a previous study (Huang et al., 2019). HEK293T cell data were generated in this study. The proportion of sites is given in parenthesis. The motifs were generated using WebLogo 3. **(B)** The numbers of NSUN2-dependent or -independent sites that are with or without a UCCA motif in HeLa and HEK293T cells. **(C)** The m^5^C level distribution in samples analyzed in **Figure 1A**. (**D-E**) Barplot showing the correlation between the expression levels of all possible methyltransferases and the number of Type II m^5^C sites in human (**D**) or mouse samples (**E**). Human and mouse gene expression data were from GTEx and modENCODE projects, respectively.

**Figure S2.**
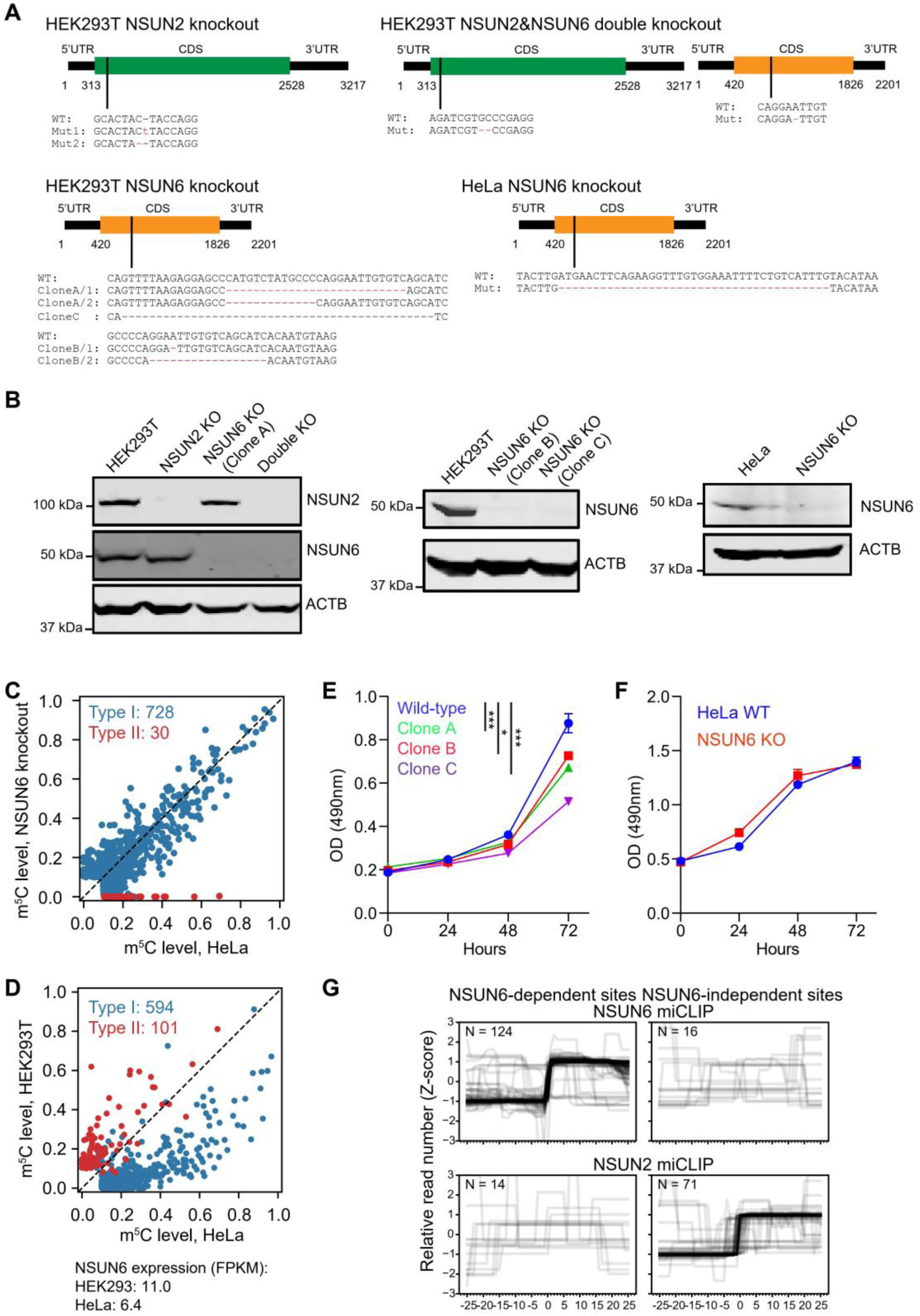
Knockout cell generation and the analysis of miCLIP data. **(A)** Schematic representation of knockout generation using the CRISPR/Cas9 system. The sequence information for the mutants is indicated. All mutant alleles result in a premature stop codon. For NSUN6 knockout HEK293T cells, we generated three independent clones (Clone A, B, and C). Clone A was used for all the experiments; Clone B and C were only used for the experiment in **Figure S2B&E**. **(B)** Western blot validation of the mutagenesis. **(C)** Comparison of methylation levels of m^5^C sites measured between wild-type and NSUN6 knockout HeLa cells. The numbers of Type II and Type I m^5^C sites are indicated. Sites covered by at least 20 reads in both samples and with a methylation level of ≥ 0.1 in either wild-type or NSUN6 knockout HeLa cells are shown. **(D)** Comparison of methylation levels of m^5^C sites measured between HEK293T and HeLa cells. The numbers of Type II and Type I m^5^C sites are indicated. Sites covered by at least 20 reads in both samples and with a methylation level of ≥ 0.1 in either HEK293T or HeLa cells are shown. **(E)** The MTS assay was used to quantify viable cells at different time points. Data are presented as Mean ± SEM (n = 4). The p values were determined using the Student’s t-test by comparing each of the mutant samples with the wild-type samples at 72 hours. *, p < 0.05; ***, p < 0.001. **(F)** The MTS assay was used to quantify viable cells at different time points. Data are presented as Mean ± SEM (n = 4). **(G)** The distributions of miCLIP reads around mRNA m^5^C sites. Normalized miCLIP read counts were plotted around the NSUN6 -dependent and -independent sites using NSUN2 and NSUN6 miCLIP data. The upstream and downstream 25 nt regions of the m^5^C sites were selected and read counts were normalized with Z-score. The number of sites (N) that can be covered by miCLIP reads is indicated.

**Figure S3.**
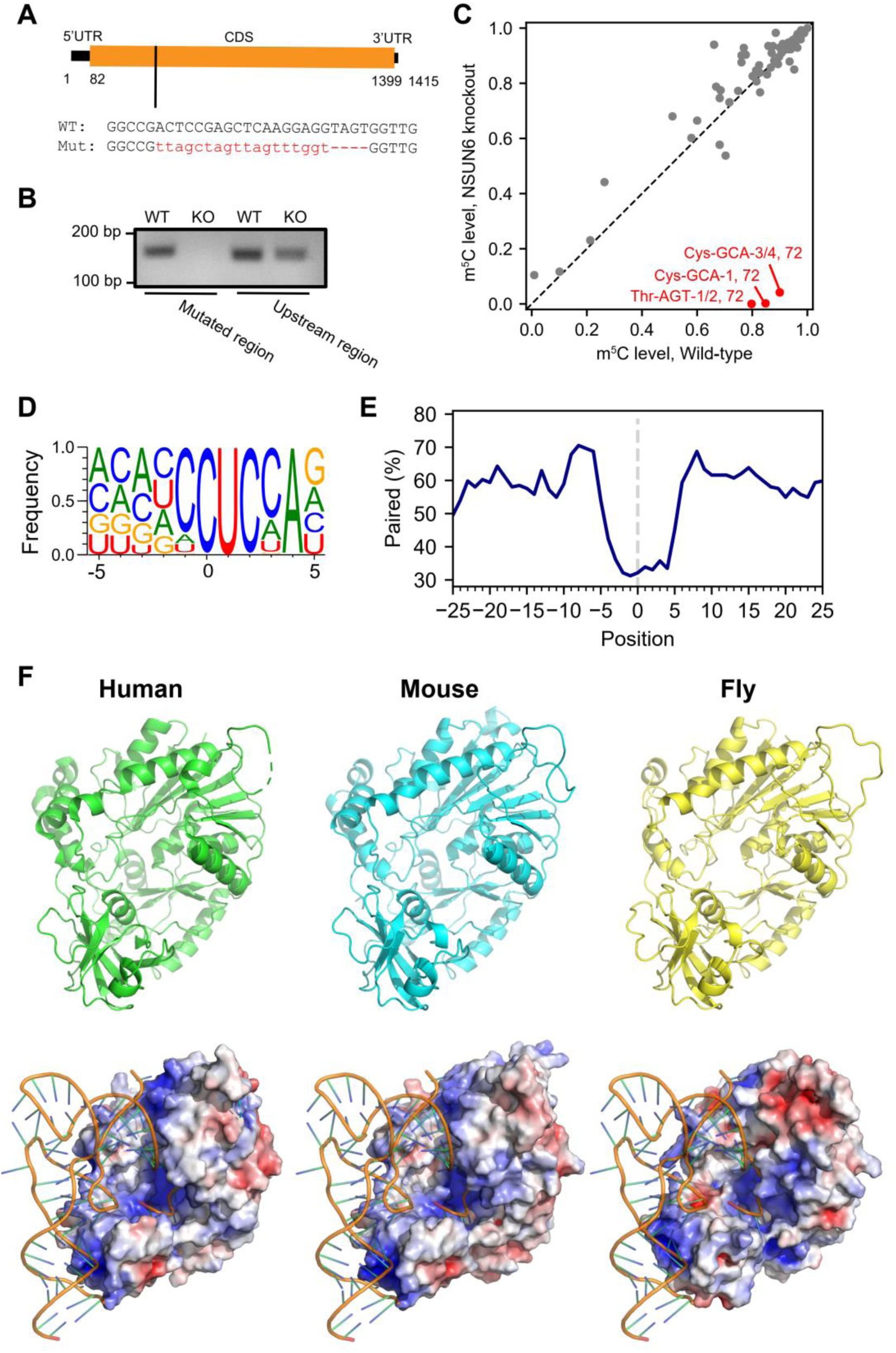
NSUN6 methylates Type II m^5^C sites in fly. **(A)** Schematic representation of NSUN6 mutant generation using the CRISPR/Cas9 system. A mutant that produced an INDEL at CDS region, leading to a premature stop codon at amino acid position 18 (the short isoform) or 99 (the long isoform), was selected. **(B)** RT-PCR verification of NSUN6 expression in wild-type and mutant flies. RNA from whole body samples of 5 days old adult male flies were used for analysis. A primer pair was designed to target the mutated region so only wild-type transcript can be amplified; another primer pair was designed to target the upstream of the mutated region so both wild-type and mutated transcripts can be amplified. **(C)** Comparison of methylation levels of tRNA m^5^C sites between wild-type and NSUN6 knockout flies. Whole body samples of 5 days old adult male flies were used for tRNA BS-seq. Sites covered by at least 10 reads in both samples and with a methylation level of ≥ 0.1 in either wild-type or mutant flies are shown. **(D)** The sequence context flanking NSUN6-dependent m^5^C sites. Adult fly ovary samples were used for mRNA BS-seq and a total of 224 NSUN6-dependent sites (methylated in wild-type (≥ 0.1) but not mutant flies (< 0.05)) were identified. **(E)** Metaprofiles of the secondary structure of NSUN6-dependent m^5^C sites and flanking regions in fly. Data were analyzed as in **Figure 1I**. **(F)** NSUN6-tRNA structure in different species. Top: the overall structure of human, mouse, and fly NSUN6 in cartoon representation. Bottom: the binding of tRNA to NSUN6 in human, mouse, and fly.

**Figure S4.**
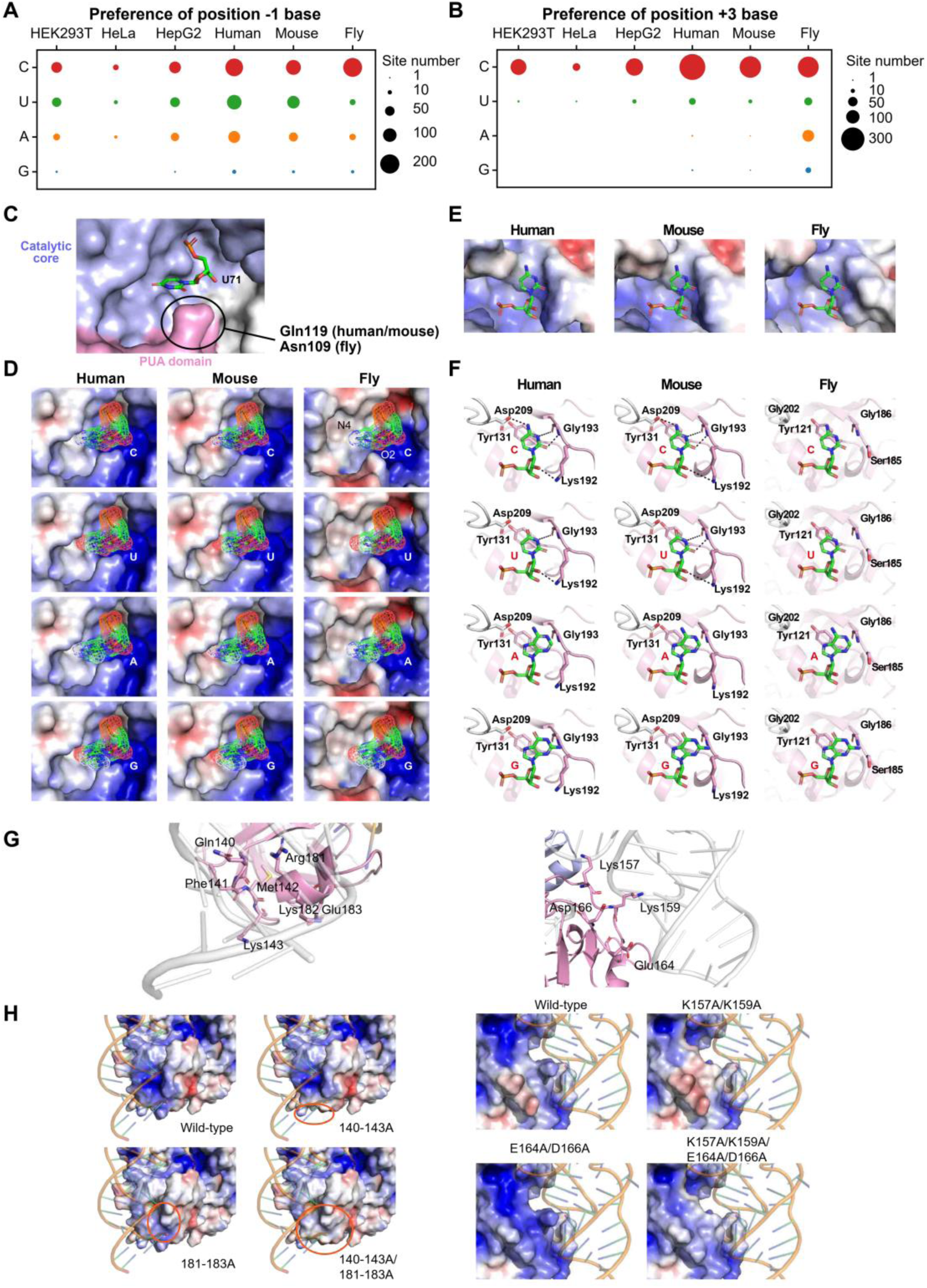
Analysis of NSUN6-RNA interactions. **(A)** The number of m^5^C sites with different base compositions at position -1. m^5^C sites identified in HEK293T cells, HeLa cells, HepG2 cells, human pooled adult tissue data, mouse pooled adult tissue data, and fly pooled sample data were analyzed. **(B)** The number of m^5^C sites with different base compositions at position +3. (**C-D**) The position -1 base and its surrounding electrostatic surface. Human NSUN6 structure was adopted from Liu et al. Mouse and fly protein structures were generated by comparative modelling. In human and mouse, steric hindrance may occur when the pyrimidines were replaced by purines (especially G), whose carbonyl group might be repulsed by the pocket. In fly, bases at position -1 are located in a more negatively charged pocket, and C may be the most favorable base in this pocket because it has a positive charge amino group towards the pocket surface and a negative charge carbonyl group towards Asn109 in fly NSUN6 (homologous to Gln119 in human). **(E)** The position +3 base and its surrounding electrostatic surface. Human NSUN6 structure was adopted from Liu et al. Mouse and fly protein structures were generated by comparative modelling. **(F)** The interactions between position +3 bases and NSUN6 residues. Possible interactions are indicated using black lines. **(G)** The residues selected to be mutated in **Figure 3A**. **(H)** The possible electrostatic surface change after replacing selected PUA domain residues by Ala.

**Figure S5.**
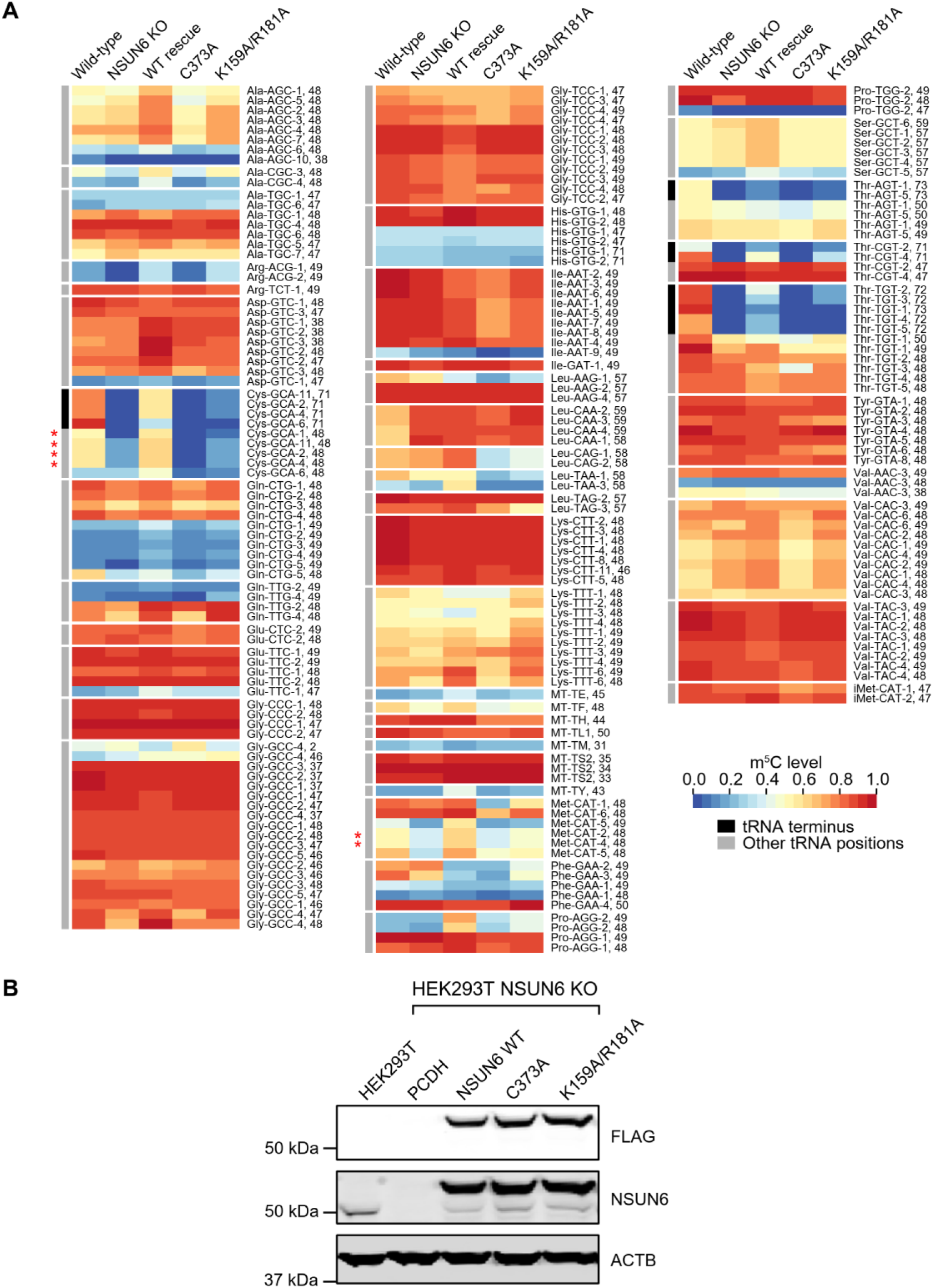
tRNA methylation profiles in different samples. **(A)** Comparison of tRNA methylation profiles between different samples. Heatmap showing the tRNA methylation profiles in wild-type cells and NSUN6 knockout cells rescued by wild-type NSUN6 or variants. Non-terminus positions that lose methylation in the NSUN6 knockout cells are marked by the asterisk. Sites covered by at least 10 reads in all samples and with a methylation level of ≥ 0.1 in wild-type cells are shown. **(B)** Western blot analysis of the NSUN6 proteins in wild-type cells or NSUN6 knockout cells overexpressed with different variants individually. ACTB served as a loading control.

**Figure S6.**
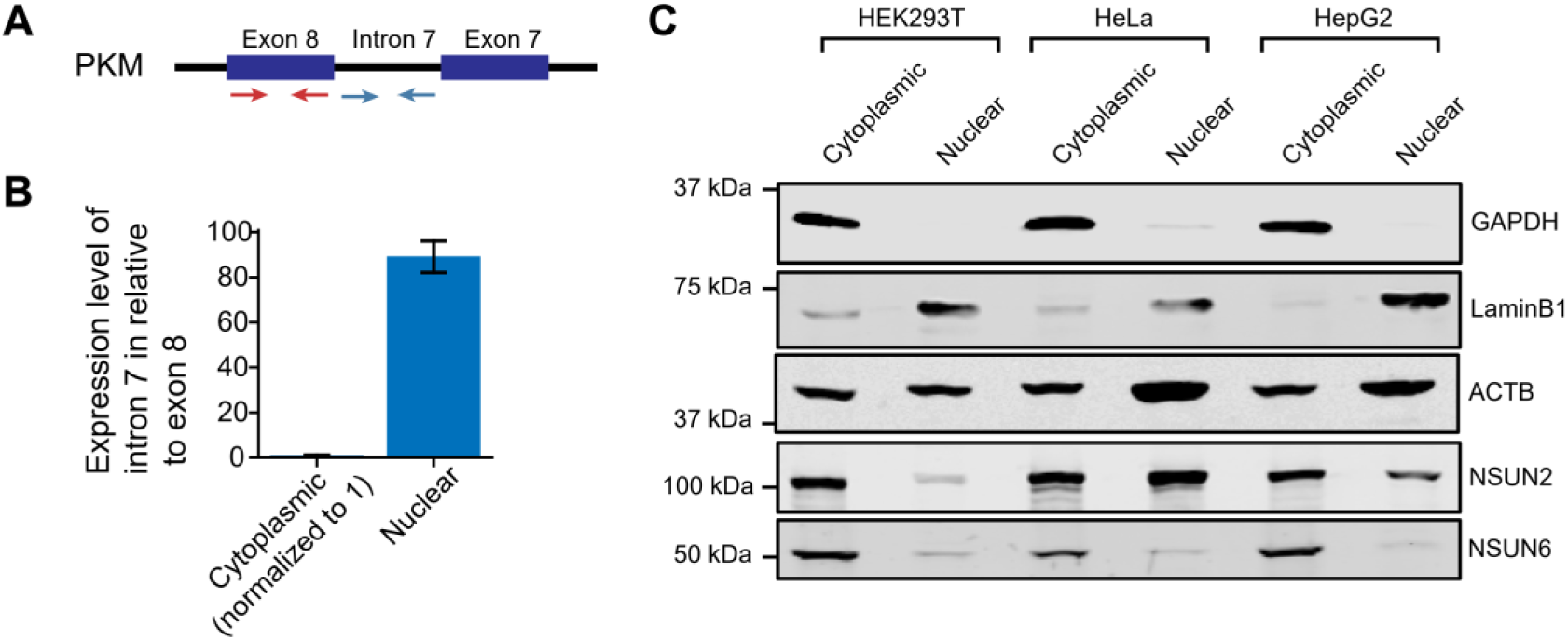
Characterizing the deposition of Type II and Type I m^5^C sites. (**A-B**) qPCR validation of cytoplasmic and nuclear RNA separation. In brief, we randomly selected a gene expressed in HEK293T cells based on ENCODE data and confirmed that intron 7 reads were only observed in nuclear RNA-seq data set (data not shown). Next, we designed two primer pairs to amplify exon 8 (red arrows) and intron 7 (blue arrows) (**A**) and performed qPCR using cytoplasmic and nuclear RNA fractions. Finally, the expression level of intron 7 was first normalized to exon 8 in each fraction, and then the relative expression of intron 7 in nuclear fraction as compared to that in cytoplasmic fraction (set to 1) was calculated. **(C)** Western blot verification of cytoplasmic and nuclear fraction separation in HEK293T cells, as well as the western blot analysis of NSUN2 and NSUN6 expression in cytoplasmic and nuclear fractions of HEK293T, HeLa and HepG2 cells. GAPDH, cytosolic marker; LaminB1, nuclear marker.

**Figure S7.**
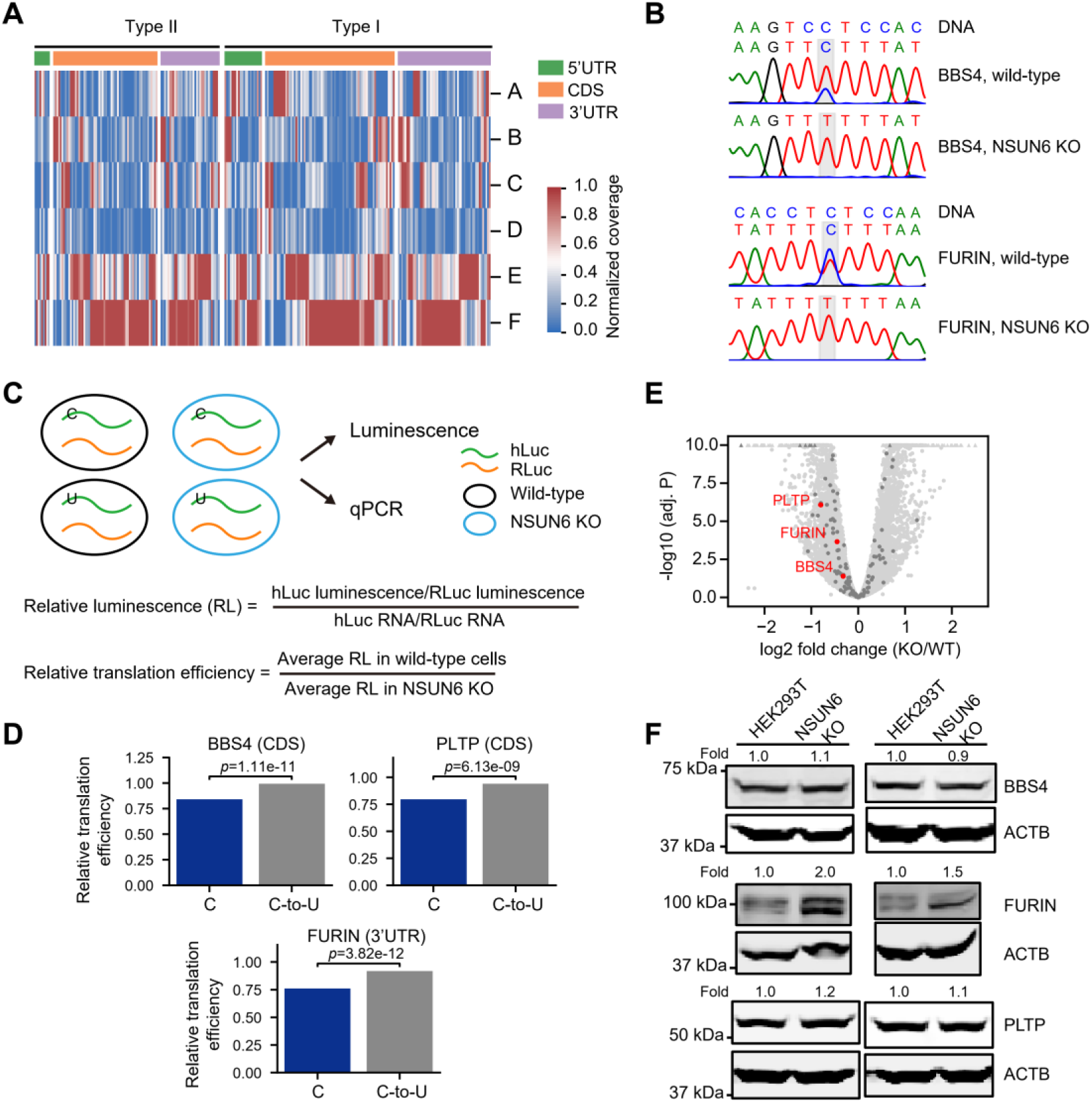
A weak negative correlation between mRNA m^5^C and translational efficiency. **(A)** Normalized read coverage of m^5^C sites across the polysomal fractions. Vertical lines represent methylated Type I or Type II m^5^C sites. The data were Min-Max normalized as in **Figure 4F**. 106 Type II and 154 Type I m^5^C sites with at least 20 reads in at least 5 fractions are shown. **(B)** BS-PCR followed by Sanger sequencing of PCR amplicons shows that the transiently expressed reporter genes were methylated. Notably, due to the presence of homopolymer (polyU, C-to-U converted RNA) in both upstream and downstream of PLTP m^5^C site, we were not able to evaluate the methylation status of this reporter gene. **(C)** Illustration of the dual-luciferase reporter assay used to examine the impact of a single m^5^C site on translational control. **(D)** Four Type II m^5^C sites (3 CDS sites and 1 3’UTR site) were examined (n=3 independent experiments). The p values were calculated using the two-way ANOVA test (P-values of F-test of base factor (C and C-to-U) in two-way ANOVA test). The source data are provided in **Table S4**. **(E)** Comparison of gene expression levels between NSUN6 knockout and control HEK293T cells via RNA-seq. Genes containing m^5^C sites selected for the reporter assay are highlighted in red. **(F)** Western blot showing the protein levels of selected genes in NSUN6 knockout and control HEK293T cells. Two biological replicates were performed.

**Note S1. The UCCA motif is a robust signature to distinguish NSUN2-dependent sites from NSUN2-independent sites**.

To utilize the UCCA motif alone to distinguish NSUN2-dependent sites from NSUN2-independent sites, it is required that NSUN2-dependent sites do not have the UCCA motif, and meanwhile, NSUN2-independent sites contain the UCCA motif. To ask whether this is the case, we analyzed mRNA BS-seq of NSUN2 knockout HeLa and HEK293T cells. We found that NSUN2-dependent and -independent sites had the G-rich triplet and UCCA motifs, respectively, in both HeLa and HEK293T cells (**Figure S1A**). Moreover, nearly no NSUN2-dependent sites had a UCCA motif and most NSUN2-independent sites contained a UCCA motif (**Figure S1B**). Thus, the UCCA motif is a robust signature for Type II m^5^C site identification with low false-assignment rate of Type I m^5^C sites.

**Note S2. Estimating the proportion of NSUN6-dependent m**^**5**^**C sites in HEK293T cells based on knockout data**.

To calculate the proportion of NSUN6-dependent m^5^C sites in HEK293T cells, we compared the methylation profiles between wild-type and NSUN6 knockout cells. NSUN6-dependent sites were defined as sites that 1) were methylated in wild-type cells (mismatch frequency ≥ 0.1, coverage of C+T ≥ 20 and p < 0.001), 2) had coverage of C+T ≥ 20 in knockout cells, and 3) had mismatch frequency < 0.05 in knockout cells. Of the 281 m^5^C sites identified in wild-type cells and with enough depth in knockout cells, 140 sites had a mismatch frequency < 0.05 in knockout cells. Thus we estimated that 49.8% of m^5^C sites were NSUN6-dependent.

**Note S3. The impact of NSUN6 on tRNA methylation**.

Position C72 of different isodecoders of tRNA^Thr-TGT^, tRNA^Thr-CGT^, tRNA^Thr-AGT^, and tRNA^Cys-GCA^ had different responses in the K159A/R181A rescue experiment (**Figure S5A**). Isodecoders such as tRNA-Cys-GCA-10 and tRNA-Cys-GCA-6, which have an identical or similar D-stem as the tRNA^Cys-GCA^ substrate used in the previous experiment (Liu et al., 2017), completely lost their methylation in K159A/R181A mutant rescue cells. However, very low-level methylation was detected in other isodecoders. This may be because other isodecoders have a subtle difference in the complex conformation, and the previous experiment only reflects the details of a subset of the isodecoders. Hence, further NSUN6 design is needed to eliminate the terminus methylation in all isodecoders. Interestingly, both increased and decreased methylation levels of non-terminus Cs were also found in the NSUN6 knockout cells (**Figure 3C** and **Figure S5A**). For example, we found that, in position C48 of tRNA^Cys-GCA^, the methylation was lost in the NSUN6 knockout cells (**Figure 3C** and **Figure S5A**). This result suggests that NSUN6 may indirectly affect the methylation of non-NSUN6-target sites, possibly by competitive inhibition of other methyltransferases or affecting the tRNA processing pathway. Further studies are required to investigate the mechanisms.

## Notes

### Competing Interest Statement

The authors have declared no competing interest.

